# Agrobacteria deploy two classes of His-Me finger superfamily nuclease effectors exerting different antibacterial capacities against specific bacterial competitors

**DOI:** 10.1101/2023.03.20.533438

**Authors:** Mary Nia M. Santos, Katherine L. Pintor, Pei-Yu Hsieh, Yee-Wai Cheung, Li-Kang Sung, Yu-Ling Shih, Erh-Min Lai

**Author notes:** Corresponding author Erh-Min Lai Institute of Plant and Microbial Biology, Academia Sinica 128 Sec. 2, Academia Road, Nankang, Taipei, Taiwan 115201. Department of Ecophysiology, Max Planck Institute for Terrestrial Microbiology, Karl-von-Frisch-Str. 10, 35043 Marburg, Germany (Katherine L. Pintor). Department of Pharmacology, University of Cambridge, United Kingdom (Pei-Yu Hsieh).

## Abstract

Type VI secretion system (T6SS) assembles into a contractile nanomachine to inject effectors across bacterial membranes for secretion. *Agrobacterium tumefaciens* species complex is a group of soil inhabitants and phytopathogens that deploys T6SS as an antibacterial weapon against bacterial competitors at both inter-species and intra-species levels. *A. tumefaciens* strain 1D1609 genome encodes one main T6SS gene cluster and four *vrgG* genes (i.e. *vgrGa-d*), each encoding a spike protein as an effector carrier. Previous study reported that *vgrGa-*associated gene 2, named as *v2a,* encodes a His-Me finger nuclease toxin (also named as HNH/ENDO VII nuclease) contributing to DNase-mediated antibacterial activity. However, the functions and roles of other putative effectors remain unknown. In this study, we identified *vgrGc-* associated gene 2 (*v2c*) that encodes another His-Me finger nuclease but with distinct SHH motif differed from AHH motif of V2a. We demonstrated that the ectopic expression of V2c caused growth inhibition, plasmid DNA degradation, and cell elongation in *Escherichia coli*. The cognate immunity protein, V3c, neutralizes the DNase activity and rescues phenotypes of the growth inhibition and cell elongation. Ectopic expression of V2c DNase-inactive variants retains the cell elongation phenotype while V2a induced cell elongation in a DNase-mediated manner. We also showed that the amino acids of conserved SHH and HNH motifs are responsible for the V2c DNase activity *in vivo* and *in vitro*. Notably, V2c also mediated the DNA degradation and cell elongation of target cell in the context of interbacterial competition. Importantly, V2a and V2c exhibit different capacities against different bacterial species and function synergistically to exert stronger antibacterial activity against the soft rot phytopathogen, *Dickeya dadantii*.

## INTRODUCTION

The type VI secretion system (T6SS) is a nanomachine used by many Gram-negative bacteria for antagonism or pathogenesis by injecting toxins into target bacterial or eukaryotic cells [1–3]. T6SS is composed of a membrane complex in connection to a baseplate, which is the docking site for polymerization of a tube surrounded by a contractile sheath [4]. The tube is a puncturing device stacked of Hcp protein hexamer tipped with the trimeric Valine-glycine repeat protein G (VgrG) spike and a PAAR repeat-containing protein which sharpens the tip [5]. When the sheath contracts, the puncturing device carrying effectors is propelled towards the target cell to deliver effectors. After firing, the sheath disassembles and the subunits of the sheath are recycled to build another machine [6].

In terms of delivery, the T6SS effectors can be classified as cargo or specialized effectors. The cargo effectors interact non-covalently with Hcp, VgrG or PAAR while in specialized effectors, the effector domain is covalently fused to Hcp, VgrG or PAAR [7]. Functionally, most of the identified effectors mediate antibacterial activities including nuclease, peptidoglycan hydrolase, lipase, phospholipase, NAD(P)+-glycohydrolase and ADP ribosyltransferase by targeting conserved cellular structures such as nucleic acids, peptidoglycan, and inner membrane as well as specific metabolites or proteins [8, 9].

To date, the characterized T6SS nuclease effectors include members of HNH, N-Tox, Tox-Rease, and PoNe superfamilies and were shown to target DNA substrates without specificity [10–16]. Thus, expression of these T6SS nuclease effectors causes DNA degradation *in vitro* and/or *in vivo*. Interestingly, the majority of T6SS DNase effectors belong to HNH/ENDO VII nuclease superfamily, also named as His-Me finger endonuclease superfamily consisting of 38 distinct families [17, 18]. These nuclease superfamily proteins harbor three conserved His-Asp-His within a ∼30-amino-acid domain formed by a β1 hairpin followed by an α-helix to form a ββα-metal topology for DNA binding and hydrolysis [17, 18]. Several T6SS effectors belonging to this HNH/His-Me superfamily have been identified, including Tse7 of *Pseudomonas aeruginosa* as Tox-GHH2 [15], Tse1 of *Aeromonas dhakensis* [13], V2a of *Agrobacterium tumefaciens* as Tox-AHH [11], and two Tox-SHH family effectors Tke4 of *Pseudomonas putida* [19] and Txe4 of fish pathogen *Pseudomonas plecoglossicida* [20]. Amino acid substitution experiments on Tse7, V2a, and Tse1 have indicated that this conserved catalytic site [A/G]HH is required for the nuclease activity and toxicity [11, 13, 15]. However, the impact of SHH and HNH motifs of the Tox-SHH family effectors in nuclease activity and T6SS toxicity have not been demonstrated.

*A. tumefaciens* is a Gram-negative Alphaproteobacteria belonging to the *Rhizobiaceae* family. It is an economically important pathogen that causes crown gall disease and a gene delivery tool with capability of transforming plant cells and fungi [21]. T6SS main gene cluster is highly conserved in the genome of *A. tumefaciens* genomospecies complex [22–24]. Besides the main cluster consisting of two divergent operons, *imp* (<u>imp</u>aired in nitrogen fixation) encoding core structural and regulatory components and *hcp* operon for effectors and effector delivery components (e.g. Hcp, VgrG, PAAR). Most *A. tumefaciens* strains also encode additional *vgrG* operons encoding effector-immunity (EI) gene pairs. *A. tumefaciens* strain 1D1609 encodes four *vgrG* genes (*vgrGa, vgrGb, vgrGc, vgrGd*) with each *vgrG* module associated with three to four associated genes, named as *<u>v</u>grG-associated* gene <u>1-4</u> in cluster *<u>a/b/c/d</u>* [11]. Each *vgrG* gene is genetically linked to a conserved chaperone/adaptor gene and different known or putative EI pairs, including three effectors with a conserved N-terminal PAAR-like DUF4150 domain but distinct C-terminal effector domains (V2a, V2c, and V2d) [11] (**Figure S1**). For *vgrGb* module, V2b does not encode an effector domain but downstream two genes encode a putative ADP-ribosylating enzyme (V3b) and a Rhs-linked effector domain (V4b), followed by a putative immunity. Previous study by deletion of single or multiple EI pairs suggested that *vgrGa-associated* effector V2a, a DNase effector harboring C-terminal Tox-AHH domain, appears to be the major antibacterial toxin but *vgrGd-associated* effector V2d also contributes to full antibacterial activity for 1D1609 against *Escherichia coli* prey [11]. The biochemical and biological functions of other putative effectors encoded in the other three orphan *vgrG* modules remain uncharacterized.

In this study, we discovered that *vgrGc-*associated effector V2c is a Tox-SHH DNase toxin and explored the rationale of having two T6SS DNase toxins in *A. tumefaciens* strain 1D1609. We show that the expression of V2c caused growth inhibition, plasmid DNA degradation and cell elongation, which the phenotypes are neutralized by co-expression of the cognate immunity protein, V3c. We also show that the SHH motif is responsible for the nuclease activity of V2c. Importantly, V2a AHH nuclease and V2c SHH nuclease exert different antibacterial capacities against specific bacterial competitors and function synergistically to exert stronger antibacterial activity against the soft rot phytopathogen, *Dickeya dadantii*. Harboring two classes of His-Me finger nuclease effectors with potential target-specific toxicity may grant 1D1609 with versatile antibacterial weapons in facing different bacterial competitors in microbial community.

## RESULTS

### V2c is a DNase belonging to the Tox-SHH clade of His-Me finger superfamily

In 1D1609, *vgrGa*-associated effector V2a appears to be the major antibacterial toxin against *Escherichia coli* prey [11]. V2c is a putative PAAR (DUF4150)-linked specialized effector (**Figure 1A**) but the presence or absence of *v2c* did not affect the antibacterial activity of 1D1609 against *E. coli* prey [11]. Aside from the N-terminal PAAR domain, V2c does not show any sequence similarity to V2a (**Figure S2**). BLAST and CDD searches did not identify known or conserved domain at its C-terminal region. However, multiple sequence alignment of V2c and V2c orthologs revealed conserved SHH and HNH motifs at the C-terminal region (**Figure 1B**). These V2c orthologs all harbor C-terminal Tox-SHH domain but their N-terminal region can be fused to other known T6SS domains such as <u>F</u>ound in type s<u>IX</u> effector (FIX), DUF4150/PAAR-like domain and Rhs core domain ([25], **Figure 1C).** Using the protein homology detection program HHpred [26], we found that the C-terminal region of V2c encodes two parallel β-strands connected by α-helix, which is signature structure in His-Me finger nuclease [17]. Consistent with the result of secondary structure prediction, 3D structure prediction using AlphaFold ([27, 28], **Figure 1D**) shows this domain consists of two antiparallel β-strands connected with an Ω loop with a His (H384) at the C-terminus of the first β-strand and followed by an α-helix, featuring a HNH motif of His-Me finger endonuclease [17, 18]. These suggest V2c belongs to the Tox-SHH (Pfam PH15652) clade of His-Me finger superfamily.

**Figure 1.**
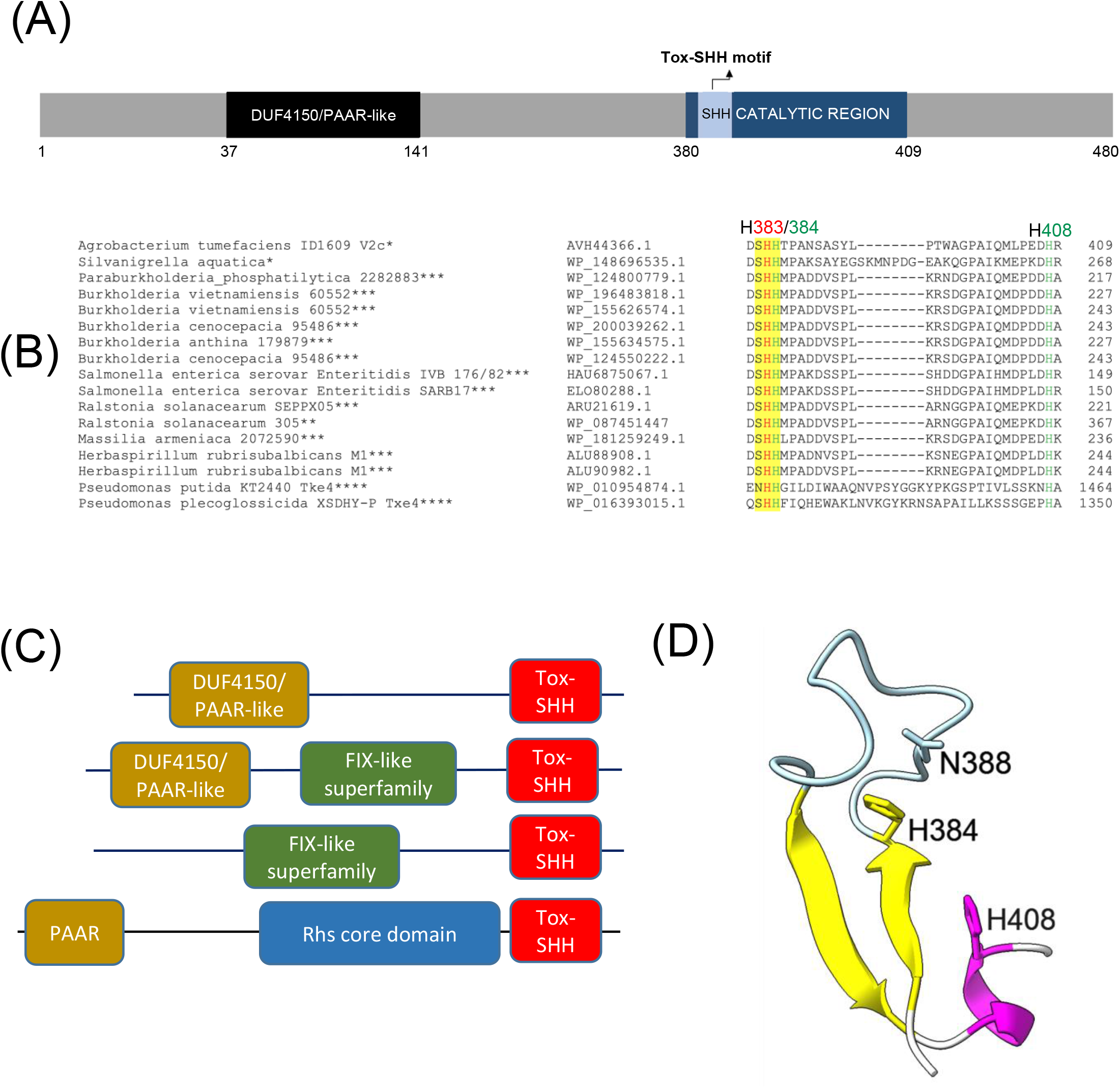
V2c is a PAAR-linked specialized effector containing C-terminal Tox-SHH motif. **(A)** Schematic diagram of V2c protein showing N-terminal DUF4150 PAAR region and the catalytic region (amino acid 380-409) which covers the Tox-SHH motif. **(B)** Partial sequence alignment of orthologs of V2c from BLASTP. The strain name and accession number are indicated on the left. SHH motif is highlighted in yellow, with H383 (indicated in red) as the metal ion coordinating residue, H384 and H408 (indicated in green) of HNH motif for catalysis and metal ion binding respectively, are targeted for mutagenesis. The star corresponds to the specific domains shown in Figure 1C: *DUF4150/PAAR-like, **DUF4150/PAAR-like + FIX-like, ***FIX-like superfamily, **** PAAR + Rhs core domain. **(C)** Domain architectures of V2c orthologs, Tox-SHH-containing proteins found in this study. Full length sequences were analyzed and the domains were from Pfam database with E value threshold set at 10^-5^. **(D)** Cartoon model of V2c (residue 380-409). The structure of V2c predicted by AlphaFold indicates V2c consisting a His-Me finger domain with two antiparallel β-strands (yellow) connected with an Ω loop (light blue), a histidine (H384) at the C-terminus of the first β-strand and followed by an α-helix (magenta). The confidence score of this region is higher than 90, indicating high structure accuracy.

To determine the antibacterial and DNase activities of V2c, we expressed *v2c* using an arabinose inducible promoter to determine if V2c is a bacterial toxin functioning as a nuclease. Growth inhibition measured by optical density was observed when *v2c* is expressed by arabinose induction in *Escherichia coli* and this can be neutralized by co-expression of its putative cognate immunity *v3c* as compared to the vector control (**Figure 2A**). Also, colony forming unit (CFU) count shows the reduced CFU of *v2c-* expressed cells and higher CFU could be recovered with co-expression of *v3c* (**Figure 2B**). In addition, plasmid degradation assay showed that the expression of *v2c* upon arabinose induction resulted in degradation of plasmid DNA in *E. coli* and this is abolished when *v3c* is co-expressed (**Figure 2C**). The data indicated that V2c exhibits DNase activity *in vivo*.

**Figure 2.**
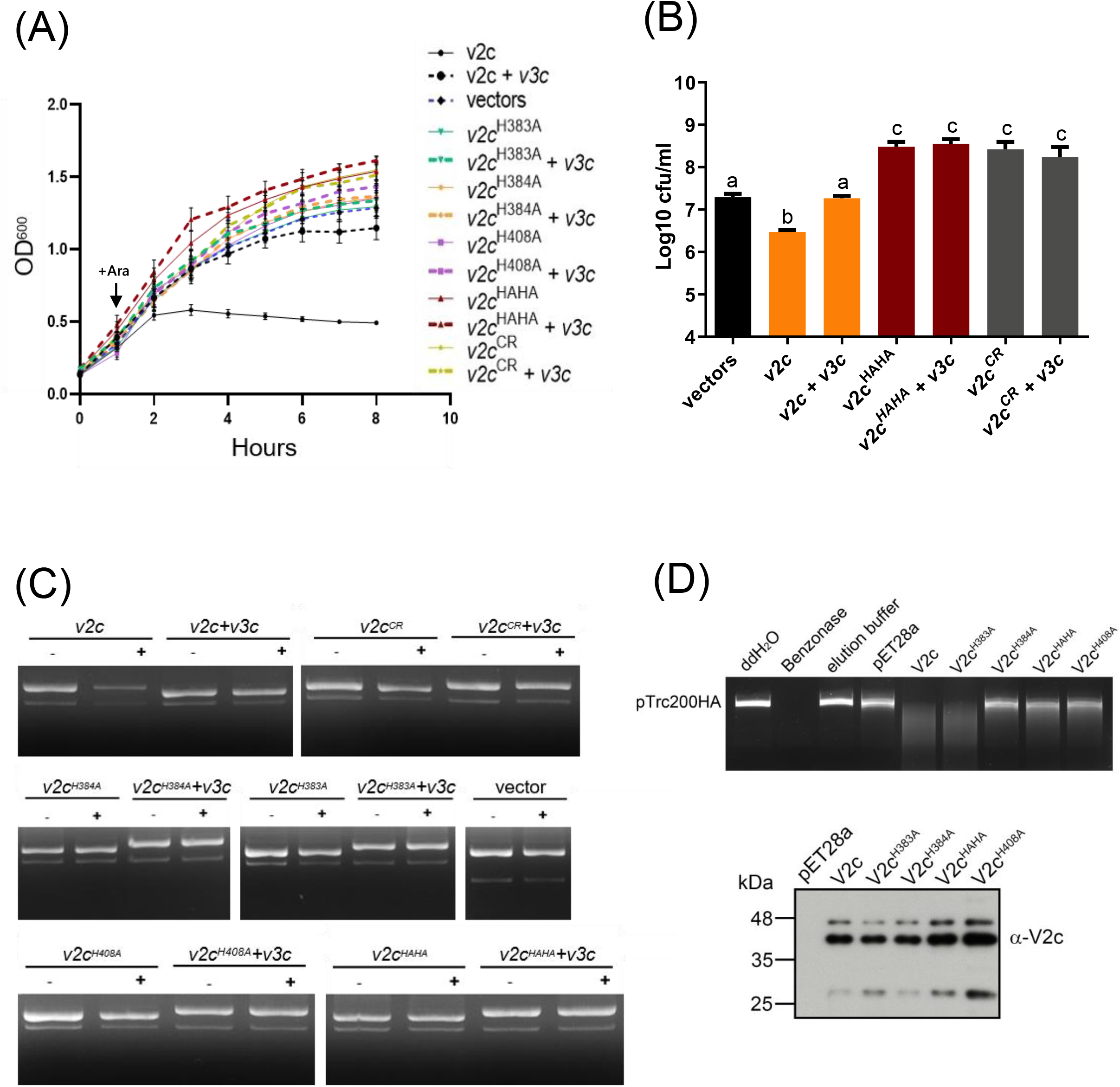
V2c exhibits growth inhibition, DNase activity *in vivo* and *in vitro.* **(A)** Growth inhibition analysis of expression of *v2c,* its variants, with or without *v3c* in *E. coli* DH10B. *E. coli* cultures were induced with 1 mM IPTG at 0 hour for *v3c* expressed from pTrc200 plasmid followed by L-arabinose (Ara) induction at 1 hour to induce *v2c* from pJN105 plasmid. Cell growth was recorded every hour at OD_600_. Strain expressing empty vectors (vectors) were used as a control. **(B)** Number of colony forming unit (log_10_ CFU/mL) quantified after growth inhibition assay at 8 hour. Data of (A) and (B) represents mean ± SEM of four independent experiments (n=4), each averaged with 3 technical repeats. Different letters above the bar indicate statistically different groups of strains (P < 0.05) determined by Tukey’s HSD test. **(C)** Plasmid degradation assay performed by *E. coli* DH10B cells harboring pTrc200 and pJN105 plasmids (vector) or the derivatives induced with (+) or without (–) Ara for 2 hour. Plasmid DNA was extracted and the degradation pattern was observed in agarose gel. **(D)** *In vitro* DNase activity assay performed by using pTrc200HA plasmid DNA as DNA substrate and incubated with purified V2c or its variants. Nuclease free water (ddH_2_O), protein elution buffer (elution buffer) and eluate from empty vector expression (pET28a) served as negative controls while Benzonase nuclease served as a positive control. The digestion of DNA substrate was determined by agarose gel electrophoresis. The reaction samples of V2c or its variants were detected by western blotting using anti-V2c antibody (α-V2c). Results of (C) and (D) are representative of three independent experiments.

We further investigated whether the SHH and HNH motifs are involved in DNase activity. His-Me finger superfamily has a strictly conserved His residue and catalytic metal ion which are both essential for nucleic acid hydrolysis. Based on this premise, we define a region spanning amino acid residues 380-409 as a His-Me finger domain core region (CR). Based on the alignment, H384 is the catalytic His and H408 is for metal ion binding while H383 may function as second residue for metal ion coordination. To verify their roles in antibacterial and DNase activity, a series of deletion as well as single and double amino acid substitution variants were generated. Strikingly, no growth inhibition and plasmid degradation were observed in any of the mutants expressing the CR deletion (V2c^CR^), single (V2c^H383A^, V2c^H384A^ and V2c^H408A^) and double (V2c^HAHA^ substitution in both H383 and H384) variants (**Figure 2A, C, D**). Higher CFU was also recovered from *E. coli* culture expressing these mutants alone or with *v3c* (**Figure 2B**). To further confirm the DNase activity of V2c, each of V2c and its variants is fused with 6x-His and purified for *in vitro* DNase activity assay. The results showed that purified V2c fraction is able to degrade DNA, but not the eluate from *E. coli* expressing the vector (pET28a) (**Figure 2D**). Consistent with the *in vivo* plasmid degradation assay, no activity could be detected from V2c^H384A^, V2c^H408A^, or V2c^HAHA^ variants. However, purified V2c^H383A^ fraction exhibited DNase activity by degrading the plasmid DNA substrate. Western blot analysis of the reaction samples shows V2c and its variants used for the assay were at similar protein levels. Taken the results of *in vivo* and *in vitro* DNA degradation assays, the conserved H384 and H408 in the HNH motif are crucial for V2c DNase activity whereas H383 in SHH motif appears to be less critical for *in vitro* but may be required for *in vivo* nuclease activity.

### Ectopic expression of V2c caused DNA degradation and cell elongation

We next examined the cell morphology and DNA integrity of *E. coli* cells expressing *v2c* and the two variants (*v2c^CR^* and *v2c^HAHA^*) that are defective in DNase activity and toxicity. Cell morphology was observed by phase contrast while bacterial cell membrane and DNA were stained by FM-4-64 and Hoechst, respectively under fluorescence microscope. Hoechst signals are significantly reduced when *v2c* is expressed by arabinose induction in *E. coli* and this can be rescued by co-expression of its cognate immunity *v3c* shown in the images and quantification (**Figure 3A, B**). Hoechst signals are diminished only in *E. coli* cells producing *v2c* but not *v2c*^HAHA^, *v2c*^CR^ and vector control, suggesting that the cellular DNA degradation is caused by V2c DNase activity, which can be neutralized by its immunity protein V3c. In addition, we also observed cell elongation in *E. coli* expressing *v2c* and co-expression of *v3c* rescues the cell elongation phenotype (**Figure 3A, C**). However, *E. coli* cells expressing *v2c*^CR^ and *v2c*^HAHA^ as well as other single amino acid substitution variants (*v2c^H383A^*, *v2c^H384A^, v2c^H408A^*) do not abolish the elongated cell phenotype (**Figure 3A, C, Figure S3A, B**).

**Figure 3.**
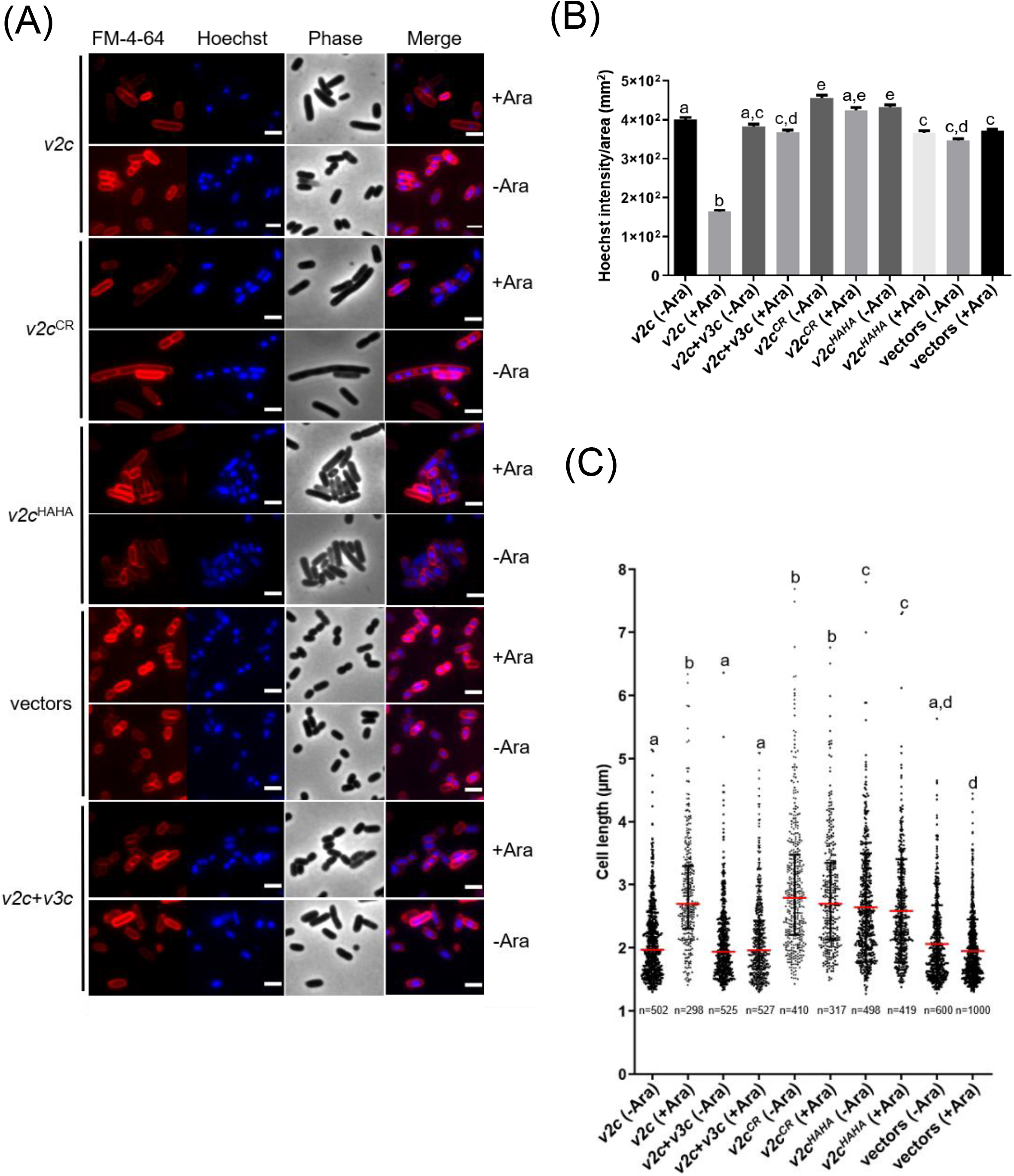
V2c Tox-SHH DNase exhibits cell elongation phenotype. **(A)** Images of *E. coli* DH10B cells harboring vector(s) and the derivatives expressing *v2c* or its variants in the presence or absence of its cognate immunity gene *v3c* with (+Ara) and without (-Ara) Arabinose induction stained with FM-4-64 (red) and Hoechst stain (blue). The micrographs from left to right are FM-4-64, Hoechst, phase contrast and a merged image of FM-4-64 and Hoechst stains. Scale bar, 2 μm. Representative images of two independent experiments are shown. **(B)** Normalized Hoechst intensity (intensity/unit area of cells) and **(C)** Cell length (μm) as measured from the indicated number of cells (n) per treatment. The graph shows a combined count from three frames of a representative result of two independent experiments. The red line in (C) shows the median with interquartile range. Statistics were performed with data mean ± SEM of three frames. Different letters above the bar indicate statistically different groups of strains (P < 0.05) determined by Tukey’s HSD test.

### Ectopic expression of V2a also caused cell elongation phenotype

Previous study showed that V2a is a Tox-AHH nuclease that serves as the major antibacterial weapon in 1D1609 [11]. H385 and H386 are two of the predicted His residues in AHH motif of V2a, corresponding to H383 and H384 in SHH motif of V2c. The structure predicted by AlphaFold indicates V2a also features a His-Me finger, having two α-helices with histidine residues (H430, H456) followed by the β-strands (**Figure 4A**). Previous findings demonstrated that the expression of *v2a* but not *v2a^H385A^*caused growth inhibition and plasmid degradation in *E. coli* [11]. To gather further insight of V2a AHH motif in the His-Me finger domain, we evaluated the growth and morphology of arabinose induced *v2a* mutants. Results show that the expression of *v2a* caused reduction in Hoechst-stained DNA signals and resulted in cell elongation (**Figure 4B, C, D**). The expression of *v2a^H385A^* has similar levels of Hoechst-stained DNA signals and cell length as compared to the vector control. These results suggest that the cell elongation phenotype observed were caused by V2a DNase activity, whereas V2c shows DNase activity-independent cell elongation. We also determined the toxicity of another putative effector V4b of 1D1609, a PAAR (DUF4150)-linked specialized effector but harboring a Rhs domain with structural similarity to the insecticidal toxin [11] (**Figure S4A**). Expression of *v4b* caused growth inhibition in *E. coli,* in which the growth inhibition can be rescued by co-expression of its putative cognate immunity *v5b* to a level similar to vector control (**Figure S4B**). However, no DNA degradation and cell elongation could be observed (**Figure S4C, D, E, F**). These results show that V4b is a bacterial toxin and V5b is the cognate immunity protein and suggest that cell elongation phenotype may be specific to toxicity of nuclease effectors.

**Figure 4.**
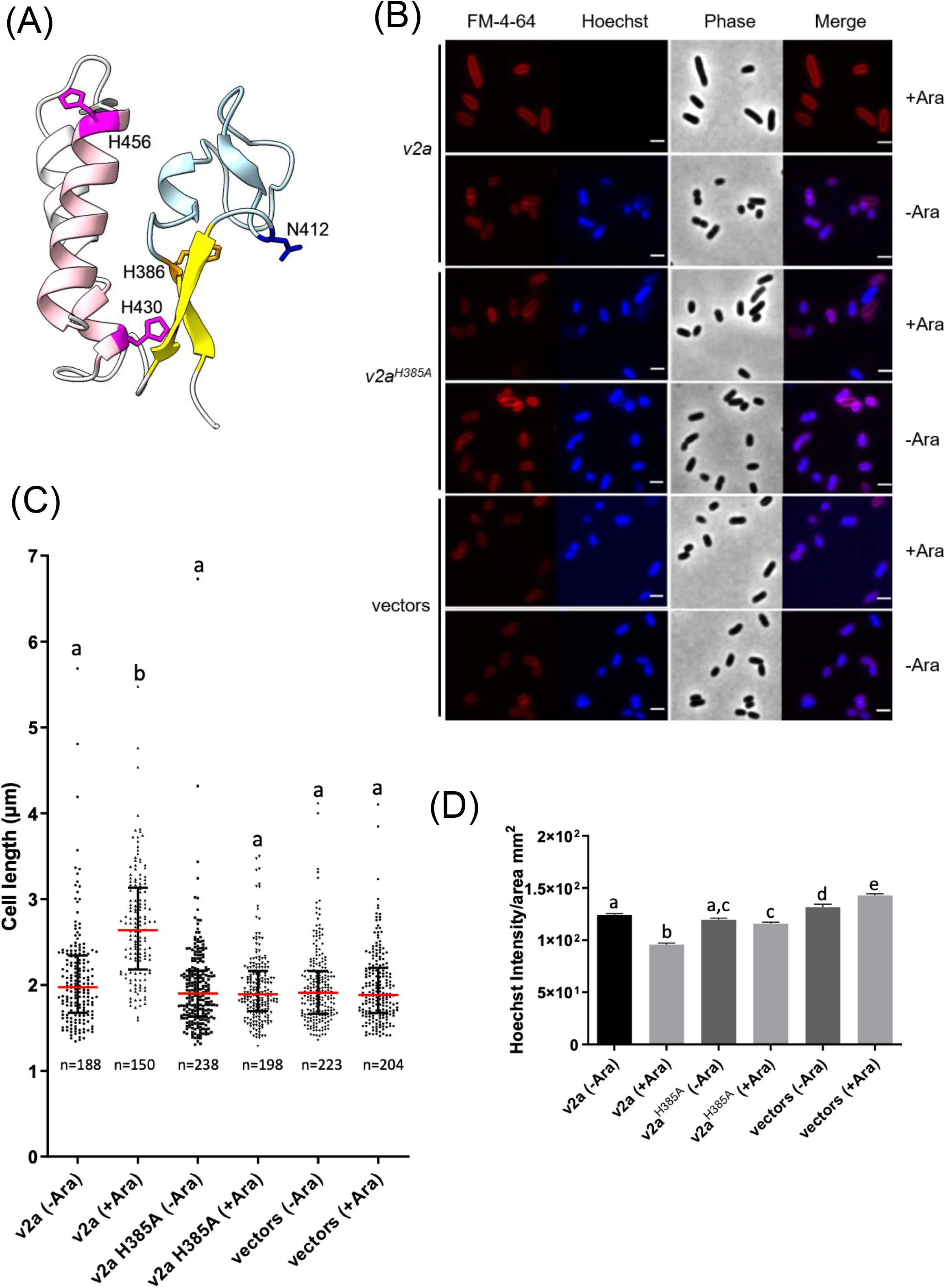
V2a Tox-AHH DNase exhibits cell elongation phenotype. **(A)** Cartoon model of V2a (residue 380-470). The structure of V2a predicted by AlphaFold indicates V2a consisting of a His-Me finger domain with its signature antiparallel β-strands (yellow) connected with an Ω loop (light blue) with a histidine (H386, orange) at the C-terminus of the first β-strand. Two α-helices (pink) with histidine residues (H430, H456 magenta) followed the β-strands. N412 is found in the loop between the β-strands and H430 is in the first α-helix (E429–G430). The predicted Local-Distance Difference Test (pLDDTs) of the His-Me domains of V2a are from confident to very high, most of the pLDDTs of the residues are > 90. **(B)** Morphological analysis of *E. coli* DH10B cells harboring vector(s) or its derivatives expressing *v2a*, catalytic site (H385A) mutant without (-Ara) or with (+Ara) arabinose induction. Cells were stained with FM-4-64 (red) and Hoechst stain (blue). The micrographs from left to right are FM-4-64, Hoechst, phase contrast and a merged image of the two fluorescent images. Scale bar, 2 μm. **(C)** Cell length (μm) and **(D)** Hoechst intensity in different treatments as determined from a combined count of three random frames of a representative result, the number of cells (n) per sample is indicated. The graph shows a combined count from three frames of a representative result of two independent experiments. The red line shows the median with interquartile range. Statistics were performed with data mean ± SEM of three frames. Different letters above the bar indicate statistically different groups of strains (P < 0.05) determined by Tukey’s HSD test.

### V2c nuclease exhibits antibacterial activity at both intra-species and inter-species competition and functions synergistically with V2a against the soft rot phytopathogen, *Dickeya dadantii*

To correlate the DNA degradation and cell elongation phenotypes observed by ectopic expression of *v2c* in *E. coli* in a more biologically relevant context, we further performed interbacterial competition assay. We first selected *A. tumefaciens* strain C58, which possess incompatible EI pairs with 1D1609 and previously demonstrated to be susceptible to 1D1609 T6SS killing [22], for intra-species competition. Each of 1D1609 mutants lacking multiple EI pairs attackers was co-cultured with C58 strain harboring pRL-GFP(S65T) prey *in vitro* on AK agar [29] and the number of viable C58 prey cells (log_10_ CFU/mL) were quantified. Similar to previous killing assay using *E. coli* prey [11], we detected ∼ 2 log of reduced prey cell survival by 1D1609 WT attacker, as compared to the *Δ4EI* mutant with deletion of all four EI pairs (**Figure 5A**). Other double *(ΔEIbd*) or triple deletion mutants (*ΔEIabd, ΔEIbcd*) exhibited compromised but detectable antibacterial activity to C58, indicating that *v2a* or *v2c* alone is sufficient to exhibit antibacterial activity to C58.

**Figure 5.**
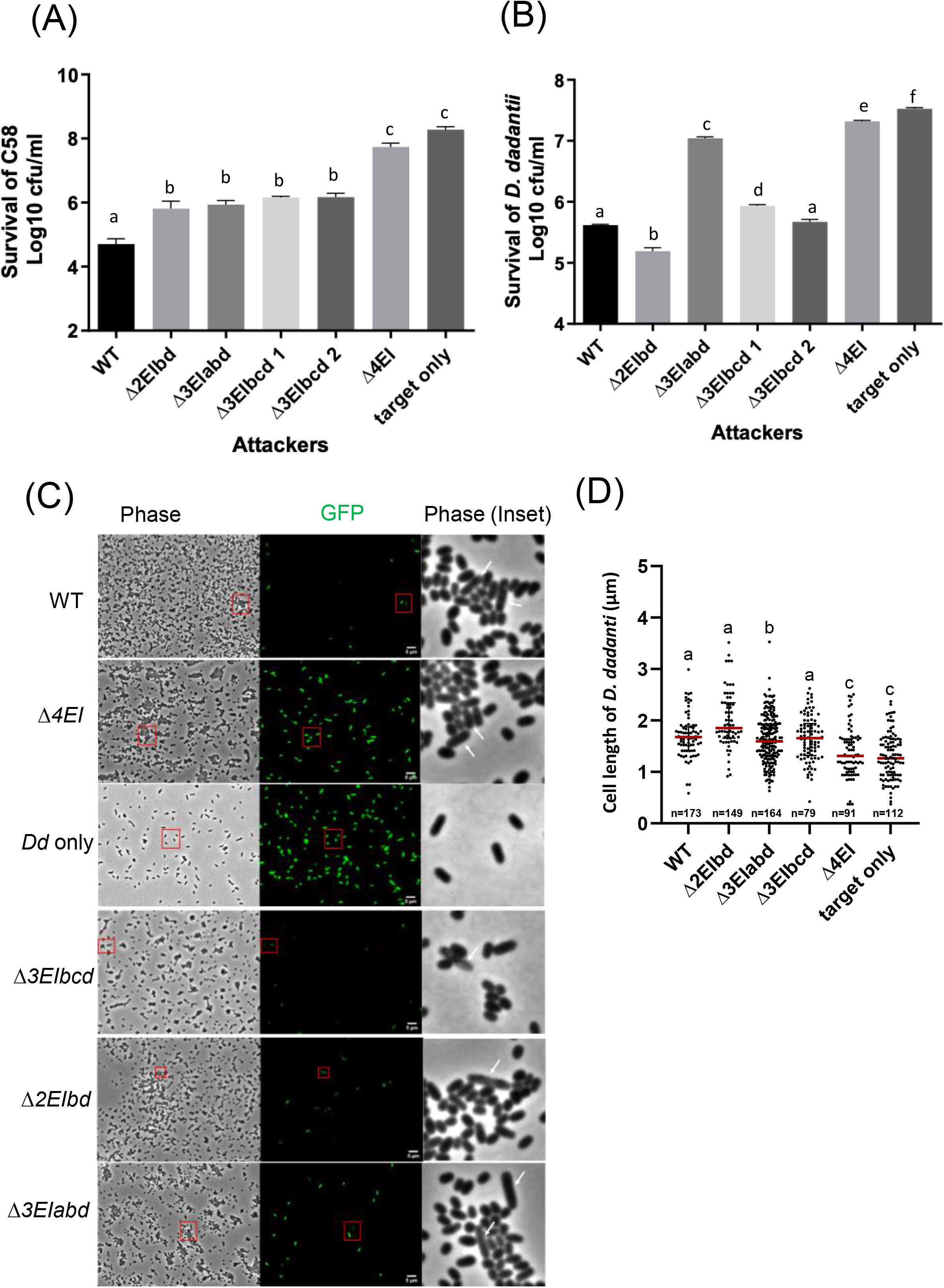
V2a and V2c exhibit different toxicity against *A. tumefaciens* and *D. dadantii*. **(A)** Intra-species competition between *A. tumefaciens* 1D1609 attacker and prey C58. **(B), (C), (D)** Inter-species competition between *A. tumefaciens* 1D1609 attacker and prey *D. dadantii* 3937 Δ*imp.* The plasmid pRL-GFP(S65T) plasmid (Gm^R^) was transformed into prey cells, which were mixed at 1:10 ratio with each of the attacker strains, 1D1609 WT or various EI mutant strains as indicated. After co-culture on AK agar plates, cells was collected, serially diluted, and plated on LB agar plate containing Gm for CFU counting of surviving prey cells **(B)**. Data are mean ± SEM (n = 6 biological repeats from three independent experiments). For imaging, co-cultured *D. dadantii* Δ*imp*-GFP(S65T) and 1D1609 EI pair mutant were concentrated and spotted on 2% agarose pad for fluorescence micrographs **(C)**. *D. dadantii* Δ*imp*-GFP(S65T) indicated as Dd only is included as a control. The inset in phase contrast shows a magnified image of cells in the red box. White arrows point to *D. dadantii* cells. Scale bar, 2 μm. **(D)** Cell length after competition between *A. tumefaciens* 1D1609 EI mutants with *D. dadantii* 3937 Δ*imp*-GFP(S65T). Cell length of GFP fluorescent *D. dadantii* cells was measured using Fiji analyze particles tool. Cell length (μm) in different treatments as determined from a combined count of at least 10 random frames of two independent experiments. The red line shows the median with interquartile range. Statistics were performed with data mean ± SEM of total counts. Different letters above the bar indicate statistically different groups of strains (P < 0.05) determined by Tukey’s HSD test.

Intrigued by the observation that *v2c* could exhibit antibacterial activity to *A. tumefaciens* C58 prey but no significant effect against *E. coli* prey [11], we reasoned that *E. coli* may not be physiologically relevant prey for *Agrobacterium* although *v2c* is able to cause toxicity when directly expressed in *E. coli* (**Figure 2A, B**). We then selected *Dickeya dadantii* as the prey cells for inter-species competition because it is a phytopathogen but also a close relative of *E. coli*, both belonging to *Enterobacteriaceae*. To avoid potential counterattack by *D. dadantii* T6SS [30], a T6SS-deficient mutant, Δ*imp* with deletion of *imp* operon, harboring pRL-GFP(S65T) (*D. dadantii Δimp*-GFP(S65T)) was used as a prey. CFU counting of *D. dadantii Δimp*-GFP(S65T) prey cells after co-culture show that ∼ 1.5 log of reduced prey cell survival by 1D1609 WT attacker, as compared to the *Δ4EI* mutant (**Figure 5B**). The presence of *v2a* only (*Δ3EIbcd*) exhibited comparable antibacterial activity as the WT while *v2c* only (*Δ3EIabd*) showed weak but detectable antibacterial activity as compared to Δ*4EI,* indicating *v2c* itself is able to kill *D. dadantii* but with much weaker activity than *v2a*. Strikingly, 1D1609 expressing both *v2a* and *v2c* (*Δ2EIbd*) could result in stronger killing effects than WT. The use of pRL-GFP(S65T) with constitutive expression of GFP also allowed us to observe the *D. dadantii Δimp*-GFP(S65T) prey cells under the fluorescence microscope after co-culture. We observed very few cells expressing GFP when competing with WT or *Δ3EIbcd* expressing *v2a* only. The lysed and elongated *D. dadantii* cells were also detected when co-cultured with 1D1609 with either strains expressing *v2a* only (*Δ3EIbcd*), *v2c* only (*Δ3EIabd*), or both *Δ3EIbd* (**Figure 5C**). We further quantified the cell length of GFP-expressing *D. dadantii* cells. The results show that majority of cells were elongated after 18 hours of co-culture with 1D1609 expressing either *v2a* (*Δ3EIbcd*) or *v2c* (*Δ3EIabd*), or combination of both (*Δ3EIbd*), as compared to those *D. dadantii* cells alone or co-incubated with *Δ4EI* (**Figure 5D**).

Taken together, these results suggest that V2c nuclease exhibits antibacterial activity at both intra-species and inter-species competition but at different capacities. While V2a and V2c exert similar antibacterial against *A. tumefaciens* C58, V2c only exhibits weak antibacterial activity but functions synergistically with V2a when competing with *D. dadantii* prey, resulting to elongated and lysed cells.

## DISCUSSION

His-Me finger nucleases are a large and diverse superfamily of nucleases; however, only a limited number of T6SS effectors have been identified as His-Me finger nuclease. In the present study, we reported that *A. tumefaciens* strain 1D1609 encodes two His-Me finger superfamily DNases, V2a and V2c belonging to Tox-AHH and Tox-SHH family respectively, exhibiting different capacities to specific bacterial competitors.

His-Me finger nucleases are one-metal-ion-dependent nucleases, primarily dependent on Mg^2+^ or in some cases dependent on Zn^2+^. The only invariant residue in His-Me endonuclease is the catalytic His located at the end of β1 strand (1^st^ H in HNH motif, H384 for V2c and H386 for V2a, **Figure 1D, 4A, S2)**, which functions as the general base for hydrolysis of phosphodiester bonds. The other conserved His (last H in HNH motif, H408 for V2c and H430 for V2a) and Asn/Asp/Glu/His located before the invariant His (H383 for V2c and H384 for V2a) are involved in coordinating the single metal ion. Indeed, our data show that H383, H384, and H408 of V2c and H385 and H386 of V2a are required for exerting DNase activity *in vivo* (**Figure 2C**, [11]. However, different from the loss of DNase activity *in vivo* and *in vitro* for both V2c^H384A^ and V2c^H3408A^, V2c^H383A^ variant retains the DNase activity by an *in vitro* nuclease activity assay (**Figure 2D**). This indicates the requirement of H383 in coordinating metal ion binding for DNase activity may be subject to environmental factors.

His-Me finger encodes a small structural motif, so it is usually associated with other domain architectures that could play additional functions. In addition, its three-element structural motif is unable to provide enough of a hydrophobic core for stability [18]. Thus, many proteins containing the His-Me finger employ a variety of additional structural elements or domains for stabilization and/or specificity. V2c and its orthologs commonly found in *Rhizobiaceae* are not included in the reported 77 sequences with conserved core regions of His-Me finger superfamily [18]. Our evidence further suggests that V2c may represent a new branch of Tox-SHH toxin family (Pfam PF15652), which includes proteins with additional domain architectures (**Figure 1C**). These additional domain(s) are conserved regions found in many bacterial toxin proteins and may facilitate the secretion of nuclease and affect its toxicity to the competing cell. While both V2c and V2a harbor the conserved N-terminal DUF4150 PAAR-like domain, they share low sequence similarity in the C-terminal His-Me finger domain (**Figure S2**), in which the protein structures predicted by AlphaFold reveal the structural difference between V2c and V2a (**Figure 1C, 4A**). This discrepancy may contribute to the difference on toxicity strength and the role of DNase on cell elongation. Alternatively, V2a and V2c may have the same mechanism to bind and hydrolyze DNA for toxicity and the difference in the phenotype is caused by another domain outside the His-Me finger fold.

Cell elongation could still be observed in the *E. coli* strain carrying different DNase defective *v2c* variants, even in the absence of arabinose (**Figure 3A, B, S2A**). Several studies reported the leaky expression of T6SS toxin driven by arabinose-inducible P_BAD_ promoter resulted in toxicity in the absence of arabinose [16, 25]. Thus, we speculated that the leaky expression of these *v2c* variants is sufficient to induce the cell elongation phenotype. The data suggest that another domain outside the SHH catalytic region in V2c is responsible for the cell elongation phenotype. This is different from the nuclease activity-dependent filamentation observed previously [15, 31] and V2a (**Figure 4**), suggesting that V2c-induced cell elongation is independent of nuclease activity. The proposed biological role of bacterial filamentation is mainly related to stress response to thrive for survival [32]. One of the conditions that lead to filamentation is DNA damage. When the DNA is damaged, the SOS response is induced and cells may delay the cell cycle and inhibit cell division until DNA repair and replication are complete. Although V2c catalytic site variants lose activity for DNA cleavage, they may still bind to DNA and induce cell elongation. Furthermore, cell division arrest could be also caused by non-nuclease effectors. For example, a T6SS ADP-ribosyltransferase effector called Tre1 in *Serratia proteamaculans* was reported to inactivate bacterial cell division Tre1 by modifying arginine on FtsZ to block polymerization [33]. Future work by domain dissection of V2c is needed to identify the domain causing cell elongation.

While HNH/His-Me T6SS nucleases are the most prevalent nuclease toxins identified in T6SS, most bacterial genomes only encode one type of HNH/His-Me nuclease effector, including Tse7 of *P. aeruginosa* [15], Tse1 of *A. dhakensis* [13], and Tke4 of *P. putida* [19]. Besides *A. tumefaciens* 1D1609 encoding Tox-AHH (V2a) and Tox-SHH (V2c) nucleases, the use of multiple HNH/His-Me nucleases as T6SS antibacterial weapons have been identified in fish pathogen *P. plecoglossicida* [20]. *P. plecoglossicida* T6SS-2 mediates interbacterial competition and encodes four putative effectors, all contain C-terminal toxin domains belonging to HNH/His-Me superfamily but distinct classes (Txe1 with a dipeptide HH motif, Txe2 as Tox-AHH, Txe3 with HNHc, and Txe4 as Tox-SHH) [20]. Among them, Txe1, Txe2, and Txe4 exhibit *in vitro* nuclease activity and toxicity when expressed in *E. coli*. However, only Txe1 and Txe4 contributes to interbacterial activity against *E. coli,* with Txe1 as the predominant toxin, in which its C-terminal conserved dipeptide HH motif is required for nuclease activity and toxicity to *E. coli.* Txe4 is also required for full interbacterial competition but the role of SHH motif in DNase and toxicity has not been determined. Together with our findings, it is possible that having multiple distinct or same classes of nuclease effectors may provide versatile toxicity when competing with different preys, in which some toxins may only exhibit activity against specific preys.

The strain 1D1609 is unique in *A. tumefaciens* species complex with multiple VgrG spikes carrying different antibacterial effectors [11]. Our findings that *v2a* and *v2c* exhibit different capacities against different preys support previous studies that some T6SS toxins can only exert toxicity in specific bacterial species [23, 34]. From the observation of the synergism of V2a and V2c against *D. dadantii,* harboring multiple T6SS effectors in 1D1609 may provide an advantage for the strain to maintain a growth competitiveness in different niches and environments.

## MATERIALS AND METHODS

### Bacterial strains and growth conditions

Bacterial strains and plasmids are listed in Table S1. *Escherichia coli* strains were grown in Luria-Bertani (LB) medium at 37°C, *A. tumefaciens* strains in 523 medium at 25°C and *D. dadantii* 3937 in LB medium at 30°C. Antibiotics were added when necessary: for *E. coli*, 30 μg/mL gentamicin and 250 μg/mL streptomycin; for *D. dadantii* and *A. tumefaciens,* 50 μg/mL gentamicin.

### Molecular cloning

Plasmid pJN105 was used for expression of toxin genes and pTrc200 plasmid was used for expression of immunity gene. PCR was amplified with KAPA HiFi HotStart DNA Polymerase (Roche, Switzerland) using genomic DNA of *A. tumefaciens* 1D1609. The primers used in the study are listed in Table S2. Site directed mutagenesis was done using either DpnI or Gibson Assembly (New England BioLabs, USA), ligated using T4 ligase (Takara) and transformed into chemically competent *E. coli* DH10B. Plasmid DNA was isolated using QIAprep miniprep kit (Qiagen, Germany). All constructs were confirmed by sequencing. The in-frame deletion mutants in *A. tumefaciens* were generated using a suicide plasmid via double crossing over by electroporation or by conjugation as described [11].

### Bioinformatic analysis

The orthologs of V2c were analyzed using amino acid sequences after the DUF4150 domain via BLASTP search. Multiple sequence alignment was performed using MUSCLE or Multiple Sequence Comparison by Log-Expectation [35]. Full length sequences of the V2c orthologs were analyzed to obtain domain architectures from Pfam database [36]. The E-value threshold was set at 10^-5^. HHpred [26] and Swiss Model [37] were used for homology-based protein structure prediction. Protein structure was predicted by AlphaFold [27, 28] from the UniProt database. Secondary structure prediction was done using default settings in PSIPRED 4.0 Protein Structure Prediction Server/DMPfold 1.0 Fast Mode [38, 39].

### Growth inhibition assay

Growth inhibition assay was performed as described previously [10]. In brief, overnight cultures of *E. coli* DH10B strain with empty vectors (pJN105 for effector gene and pTrc200 for immunity gene) or the derivatives were adjusted to OD_600_ of 0.1 in LB medium. 1 mM isopropyl β-D-1-thiogalactopyranoside (IPTG, Amresco) was used to induce the expression of the putative immunity protein. After 1 hour of IPTG induction, L-arabinose (Ara, Sigma-Aldrich) was added to 0.2% final concentration, to induce expression of the toxin. Cell growth was recorded every hour at OD_600_. Empty vectors were used as controls. After 8 hours, cells were harvested to quantify the number of colony forming units (log_10_ CFU/mL) by automatic diluter and plater, easySpiral Dilute (Interscience, France).

### Plasmid DNA degradation analysis in *E. coli* cells

*In vivo* plasmid DNA degradation analysis was performed as described previously [10]. In brief, overnight cultures of *E. coli* DH10B strain with empty vectors (pJN105 for effector gene and pTrc200 for immunity gene) or the derivatives grown in LB broth were harvested and adjusted to OD_600_ ∼0.3 in LB medium. *E. coli* cultures were induced with 1 mM IPTG at 0 hour for *v3c* expressed from pTrc200 plasmid followed by L-arabinose (0.2%, final concentration) induction at 1 hour to induce *v2c* from pJN105 plasmid and cultured for 2 hours. Equal amounts of cells were used for plasmid DNA extraction and equal volume of extracted DNA was resolved in agarose gel followed by ethidium bromide staining and visualized using Gel Doc XR+ UV Gel Documentation Molecular Imager Universal Hood II (Bio-Rad, USA).

### Cloning, expression and purification of V2c and mutants

For *in vitro DNase* activity assay, the effector gene *v2c* was constructed in pET28a plasmid and immunity gene *v3c* was constructed in pTrc200 plasmid. Site-directed mutagenesis was done by using overlap extension PCR. Plasmid for expressing effector protein, *v2c* or its variants, and plasmid for expressing immunity protein, *v3c*, were transformed into *E. coli* BL21(DE3) for IPTG-induced co-expression. Bacterial culture was incubated at 37 °C until OD_600_ reached 0.5, then 0.1 mM IPTG was added followed by 5-h expression at 37°C. Pellets were collected, sonicated, and expressed proteins were purified by using Ni Sepharose 6 Fast Flow histidine-tagged protein purification resin (Cytiva, Germany). Briefly, cells were lysed in extraction buffer (50 mM Tris-Cl, pH7.5, 0.1 M NaCl, 20 mM imidazole) and proteins were eluted by using elution buffer with up to 250 mM imidazole in 50 mM Tris-Cl, pH7.5, 0.1 M NaCl.

### *In vitro* DNase activity assay

A 10 µL reaction containing 200 ng of pTrc200HA plasmid DNA and 1× CutSmart Buffer (New England BioLabs, USA) with partially purified V2c or its variants normalized by the major V2c protein band was incubated at 37°C for 1 hour. The digestion of DNA was determined by agarose gel electrophoresis and visualized by SYBR Safe DNA gel stain (Invitrogen, USA). The reaction samples of V2c or its variants were scaled up for Western blotting analysis. Anti-V2c antibody was generated in rabbits against two V2c peptides, PFYWDFPNSQVGRD and KRIAMLRGDPTRYD (indicated in Figure S1) by Yao-Hong Biotechnology Inc. (Taiwan).

### Western blot analysis

Western blot analysis was performed as described [40]. In general, proteins were resolved by SDS-PAGE and transferred onto a PVDF membrane by using a transfer apparatus (Bio-Rad, USA). The membrane was probed with primary antibody against V2c (1:4000), Hcp (1:2500), TssB (1:4000) and GFP (1:1000), followed by incubation with horseradish peroxidase-conjugated anti-rabbit secondary antibody (Chemichem) (1:25000) and visualized with the ECL system (Perkin-Elmer Life Science, Boston, USA).

### Fluorescence microscopy and image analysis

*E. coli* DH10B carrying the empty vectors (pJN105 for effector gene and pTrc200 for immunity gene) or the derivatives were grown overnight at 37°C. Overnight cultures were sub-cultured with OD_600_ ∼0.01. After 1 hour, L-arabinose (Ara) was added (0.2% final concentration) to induction. Cultures were harvested after an hour and resuspended with 30 μL PBS with 0.5 % Tween 20 (PBST). Three μL of cells was placed on 2% agarose pad and imaged. To check the DNA degradation, the same protocol was followed and live cells were stained with 2 μg/mL Hoechst (Life Technologies, USA) and 1 μg/mL FM4-64FX, fixable analog of FM-4-64 (Invitrogen, USA) for 5 mins and washed with PBS to remove the excess dye.

For co-culture of *D. dadantii* Δ*imp*-GFP(S65T) with various 1D1609 EI pair mutants, strains were mixed at 1:10 ratio on AK agar, *Agrobacterium* kill-triggering (AK) medium (3 g K_2_HPO_4_, 1 g NaH_2_PO_4_, 1 g NH_4_Cl, 0.15 g KCl, 9.76 g MES in 900 mL ddH_2_0, pH 5.5) [29] solidified by 2% (w/v) agar. Cells from 16 hour co-culture at 25°C were concentrated to OD_600_ ∼10 and spotted on 2% agarose pad for imaging.

Imaging was performed using an inverted fluorescent microscope (IX81, Olympus Japan) with objective lens UPLSAPO 100XO (Olympus, Tokyo, Japan), DAPI (49000; Chroma), GFP (49002; Chroma) and Cy3 (49005; Chroma) filter set and CCD camera (ORCA-R2; Hamamatsu, Japan). The images were acquired using Olympus cellSens Dimension software.

All the image analysis was done using Fiji analyze particles tool [41]. The micrograph of FM-4-64 fluorescence was used to segment surfaces (i.e. individual bacterial cell) with background subtraction and thresholding. Feret’s diameter was used. The cell length (μm) was determined from cumulative counts of three random frames per sample. For quantification of cell degradation, the Hoechst intensity was normalized using the formula intensity/unit area of cells, where intensity is major x minor x 3.14 (π).

### Interbacterial competition assay

Bacterial competitions were carried out as described [22] with modifications. Briefly, overnight culture of *A. tumefaciens* and *D. dadantii* 3937 Δ*imp* were mixed at a 10:1 attacker to prey ratio and spotted on Agrobacterium kill-triggering (AK) agar plates [29]. The prey *A. tumefaciens* C58 or *D. dadantii* 3937 Δ*imp* was transformed with pRL662-GFP(S65T) for gentamycin selection or microscopy imaging. After 18 hour incubation at 25°C, the co-cultured cells were collected, serially diluted and plated on LB agar containing gentamicin to quantify surviving prey cells by counting CFU.

### Statistical analysis

The statistical analyses were done using GraphPad Prism version 6.01 for Windows, GraphPad Software, La Jolla California USA (www.graphpad.com). The number of technical replicates and independent biological replicates, p values and statistical tests performed are indicated in the figure legends. The mean of independent replicates was compared using the following statistical tests: Student’s t test, one-way ANOVA followed by Tukey’s multiple comparisons tests (http://astatsa.com/OneWay_Anova_with_TukeyHSD/). The error bars indicate the standard error of the mean (SEM).

## ACKNOWLEDGEMENTS

The authors acknowledge Ching-Hong Yang for providing *Dickeya dadantii* 3937 strains, which were imported under the permit 106-B-003 issued by the Council of Agriculture of Taiwan. We would like to thank the former and current Lai lab members for their help and fruitful discussion throughout this study, as well as See-Yeun Ting and Mao-Sen Liu for critically reading the manuscript and their valuable comments. We also thank Venus Marie Gaela and Thomas Boudier for the guidance and help in processing the images using Fiji and the Sanger DNA sequencing service provided by Genomic Technology Core located at the Institute of Plant and Microbial Biology, Academia Sinica. The funding was supported by grants National Science and Technology Council (NSTC) of Taiwan (107-2311-B-001-019-MY3) and Academia Sinica Investigator Award (AS-IA-107-L01) to EML, and NSTC grant (110-2311-B001-011) to YLS. YWC was supported by the postdoctoral fellowship (NSTC 110-2811-B-001-504). The funders had no role in study design, data collection and interpretation, or the decision to submit the work for publication.

## AUTHOR CONTRIBUTION

Conceptualization: MNMS, EML; Investigation: MNMS, KP, PYH, YWC, LKS, YLS, EML; Project Funding acquisition and administration: EML, YLS; Supervision: EML, YLS; Writing of original draft: MNMS, EML; Writing – methodology, review & editing: MNMS, KP, PYH, YWC, LKS, YLS, EML.

---

**Figure S1.**
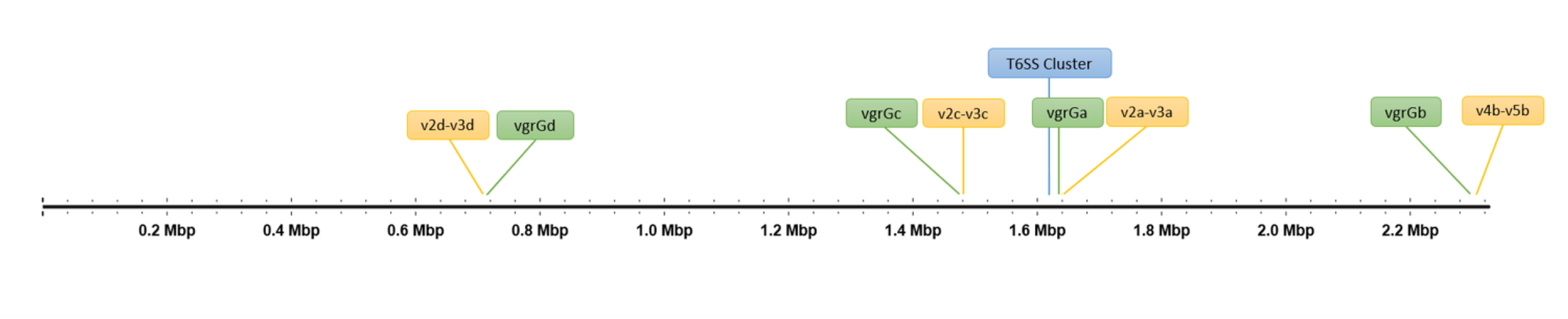
Genetic loci of *A. tumefaciens* 1D1609 T6SS gene cluster and *vgrG* genetic modules. The four *vgrG* genes (*vgrGa, vgrGb, vgrGc, vgrGd*) and their associated effector-immunity pairs are indicated on linear chromosome. The number of *vgrG-*associated genes *(v)* represents the position to *vgrG* as defined previously [11].

**Figure S2.**
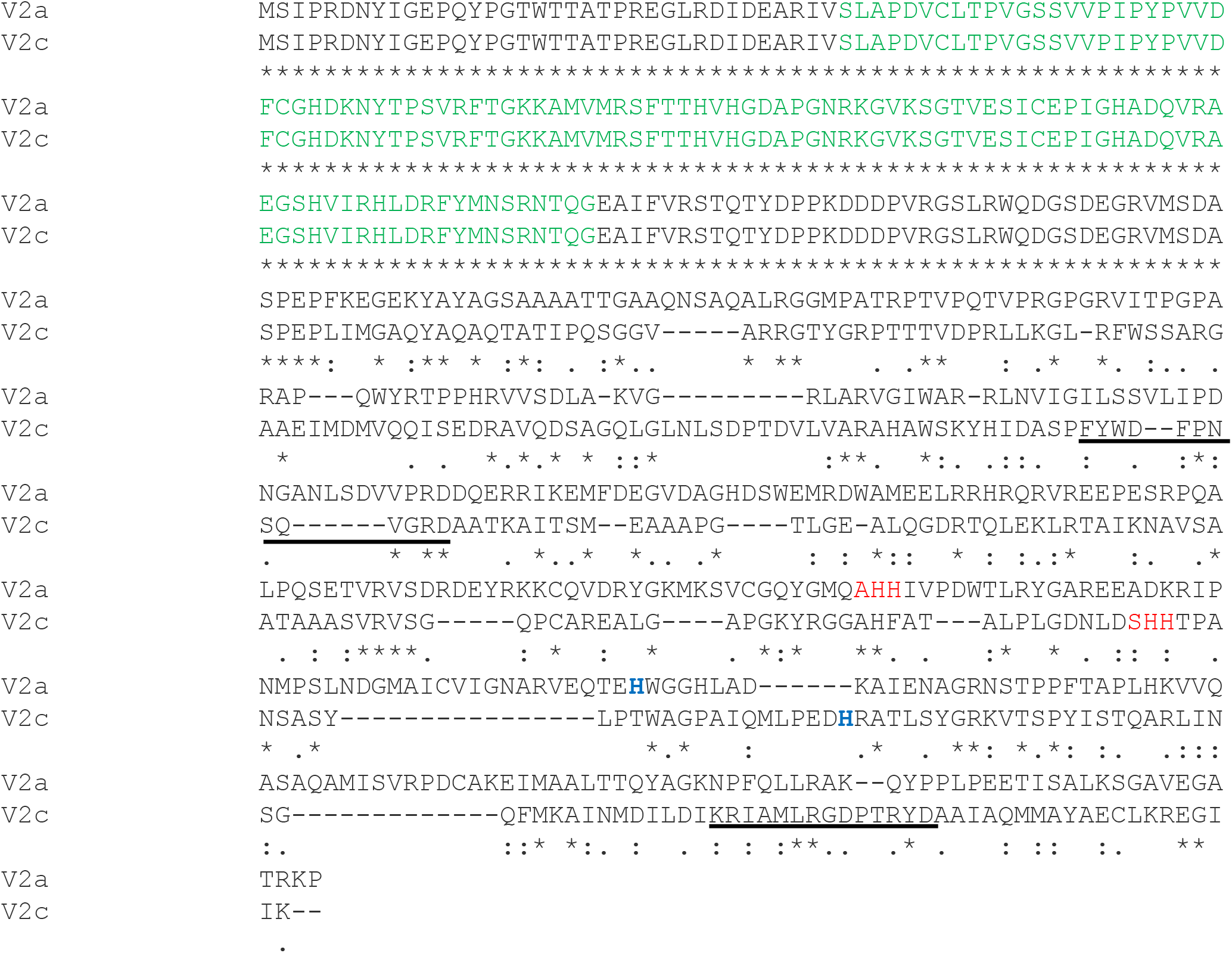
Sequence alignment of V2a and V2c. The conserved amino acid residues are indicated as ‘∗’ (identical), ‘:’ (similar) and ‘.’ (slightly similar). The DUF4150, PAAR-like domain is in green. The histidine residues in AHH and SHH motifs in His-Me finger domain of V2a and V2c respectively, V2aH385/V2cH383 and V2aH386/V2cH384, are in red. The histidine residues that predicted to responsible for metal binding in His-Me finger domain, V2aH430/V2cH408, are in a bold blue font. Two peptides used for generating antibodies were underlined.

**Figure S3.**
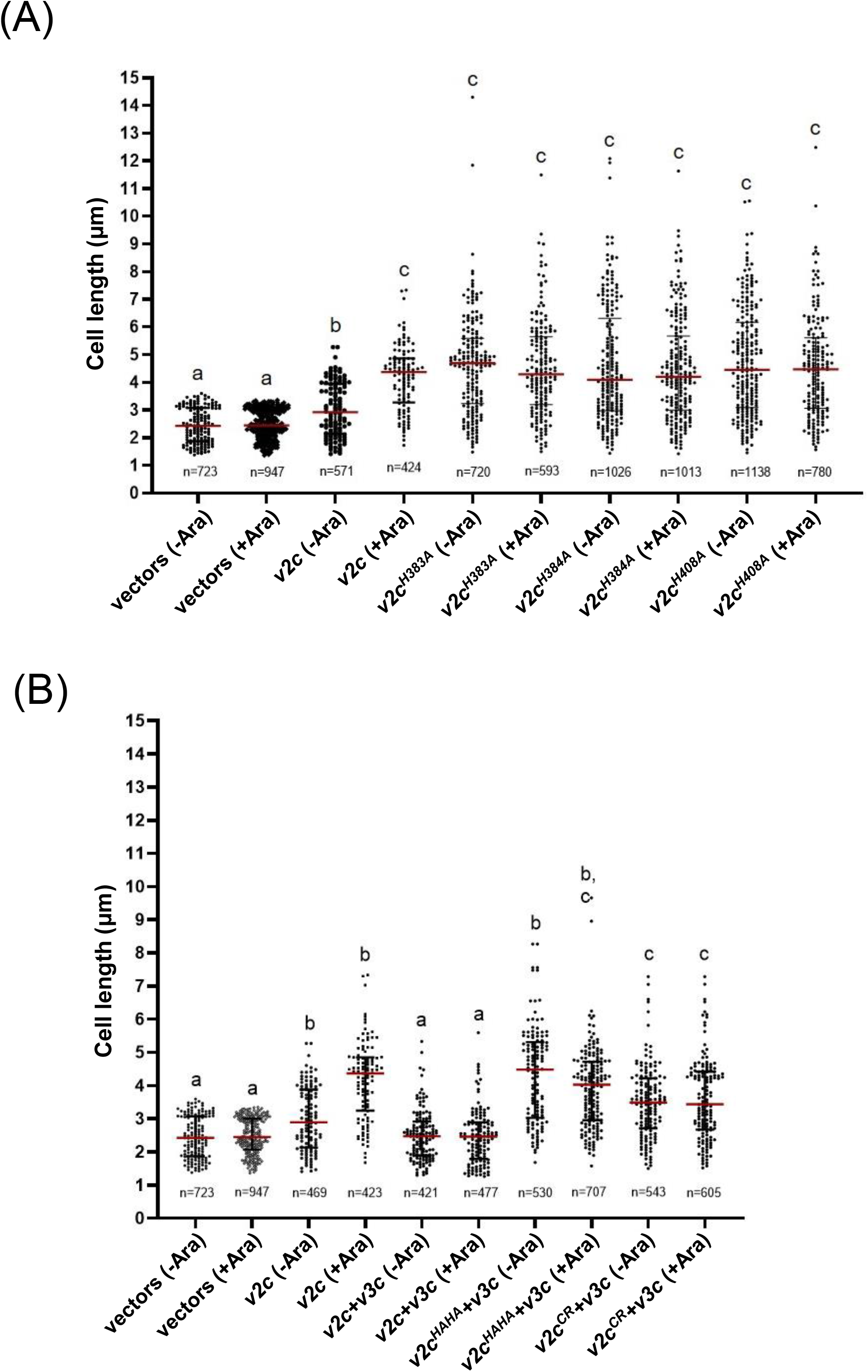
Cell length analysis of *E. coli* DH10B cells expressing *v2c* catalytic site variants. **(A)** Expression of *v2c* single catalytic site variants. **(B)** Expression of catalytic site variants co-expressed with *v3c*. Cell length (μm) as measured from the indicated number of cells (n) per sample. The graph shows a combined count from six random frames of a representative result, the number of cells (n) per sample is indicated. The graph shows a combined count from 6 random frames, 3 from each of two independent experiments. The red line shows the median with interquartile range. Statistics were performed with data mean ± SEM of 6 frames. Different letters above the bar indicate statistically different group of strains (P < 0.05) determined by Tukey’s HSD test. The red line shows the median with interquartile range.

**Figure S4.**
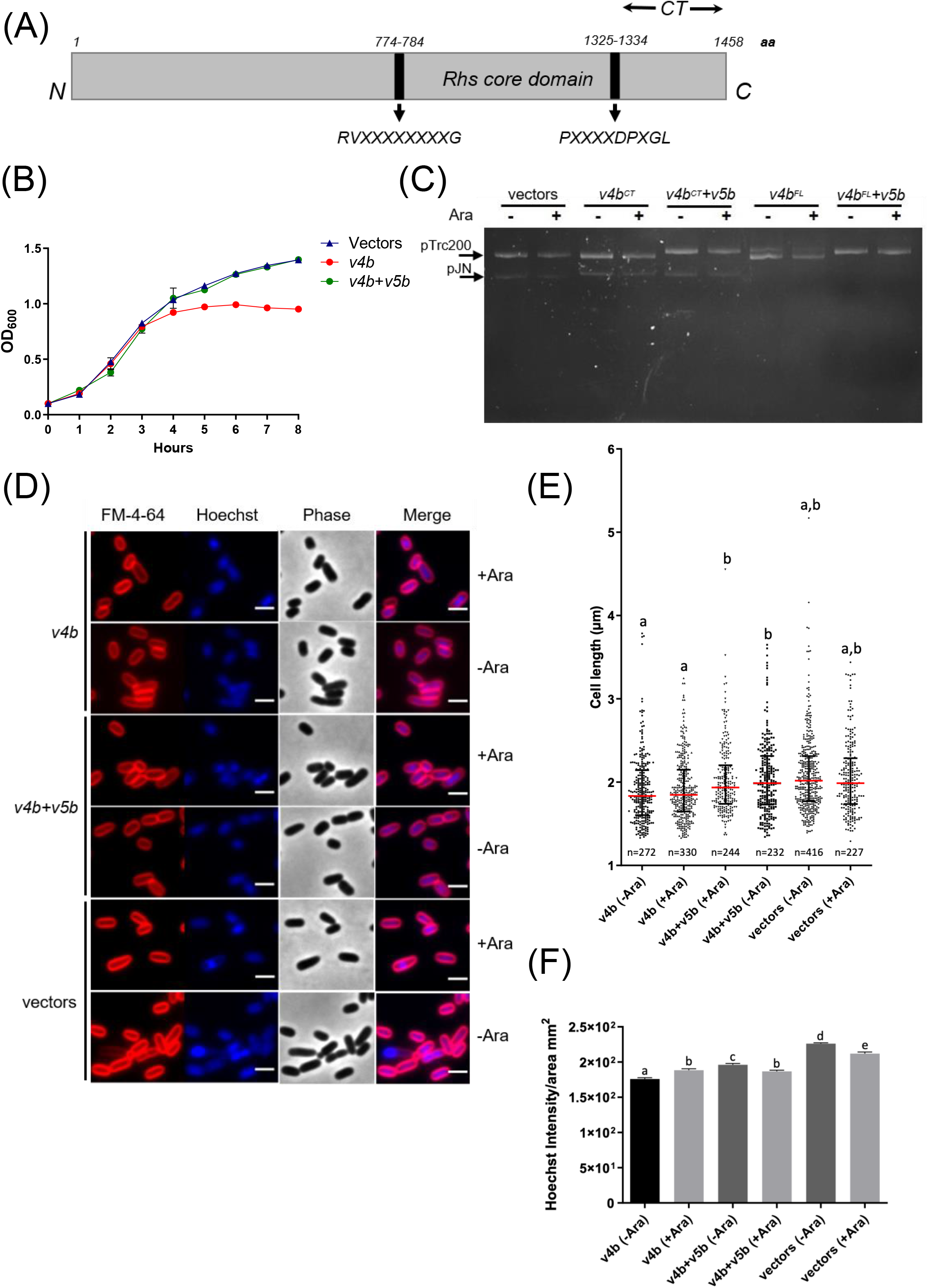
V4b exhibits growth inhibition, which could be neutralized by V5b. **(A)** Schematic diagram of V4b. The Rhs domain is bordered by conserved motifs. The CT is the C-terminal domain encoding a putative muramidase toxin. **(B)** *E. coli* growth inhibition analysis. *E. coli* DH10B cultures were induced with 1 mM IPTG at 0 hour for the expression of putative immunity gene *v5b* from pTrc200 plasmid followed by L-arabinose (Ara) induction at 1 hour to induce putative *v4b* toxin gene from pJN105 plasmid. Cell growth was recorded every hour at OD_600_. DH10B expressing empty vectors were used as a control. Data are mean ± SD of three biological replicates. Similar results were observed in three independent experiments. **(C)** Plasmid degradation assay performed by using full length V4b (*v4b*^FL^) and C-terminal domain only (*v4b*^CT^). *E. coli* cells with pTrc200 and pJN105 plasmid (vectors) or expressing *v4b* and *v4b*+*v5b* were induced with (+) or without (-) arabinose (Ara) for 2 h. Plasmid DNA was extracted and the degradation pattern was observed in agarose gel. Result shown is a representative data from two independent experiments. **(D)** Images of *E. coli* DH10B cells expressing *v4b* and *v4b* co-expressed with its cognate immunity gene *v5b* (*v4b*+*v5b*) under condition with (+Ara) or without (-Ara) arabinose induction. *E. coli* DH10B cells expressing empty vectors (vectors) under condition with (+Ara) and without (-Ara) arabinose induction served as negative controls. Cells were stained with FM-4-64 (red) and Hoechst stain (blue). The micrographs from left to right are FM-4-64, Hoechst, phase contrast and a merged image of the two fluorescent micrographs. Representative images of two independent experiments are shown. Scale bar, 2 μm. **(D)** Cell length (μm) and **(F)** Hoechst intensity in cells analyzed in (D). The red line shows the median with interquartile range. The graph shows a combined count from 3 random frames of a representative result, the number of cells (n) per sample is indicated Statistics were performed with data mean ± SEM of three frames. Different letters above the bar indicate statistically different groups of strains (P < 0.05) determined by Tukey’s HSD test.

**Supplementary Table S1.**
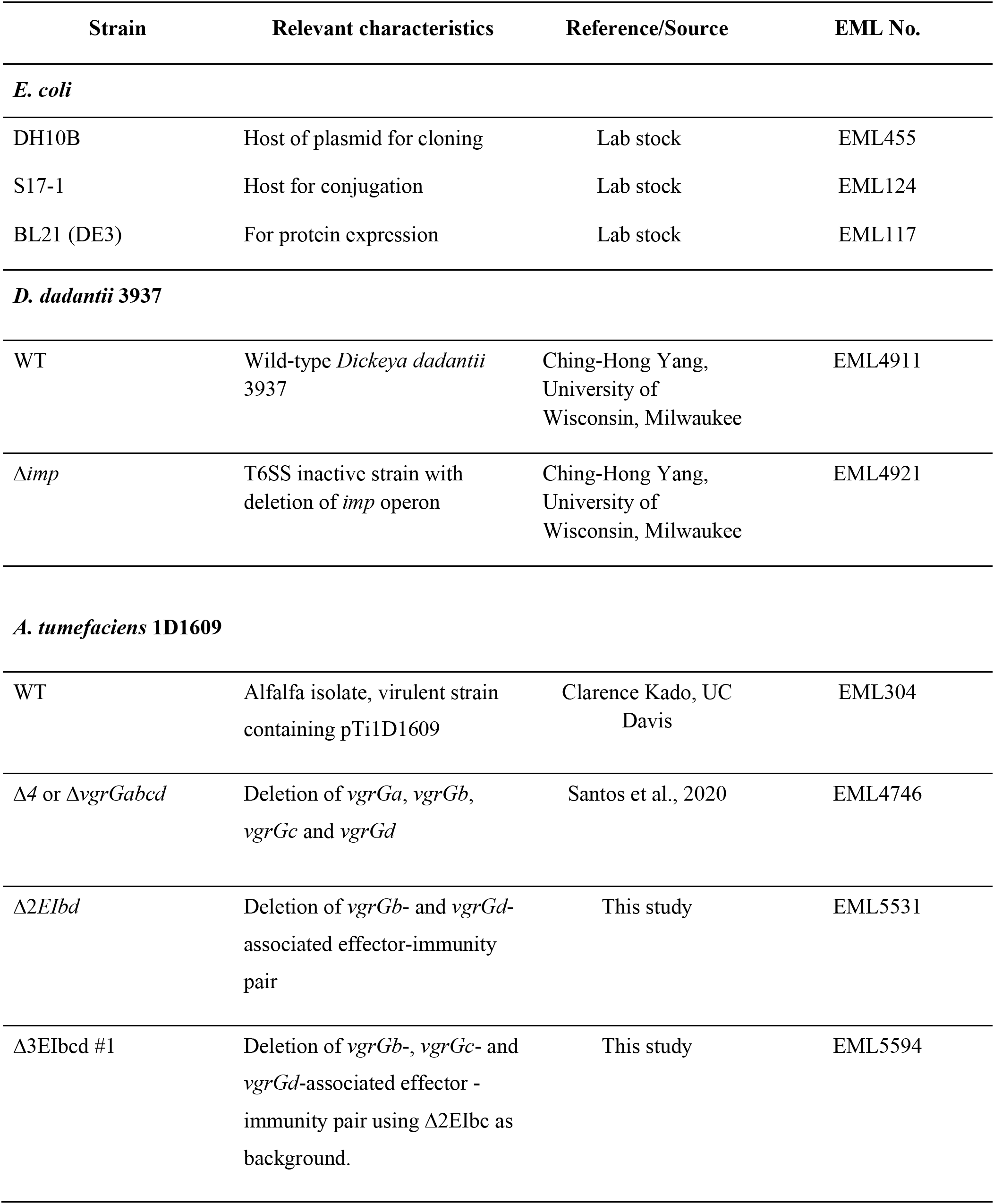
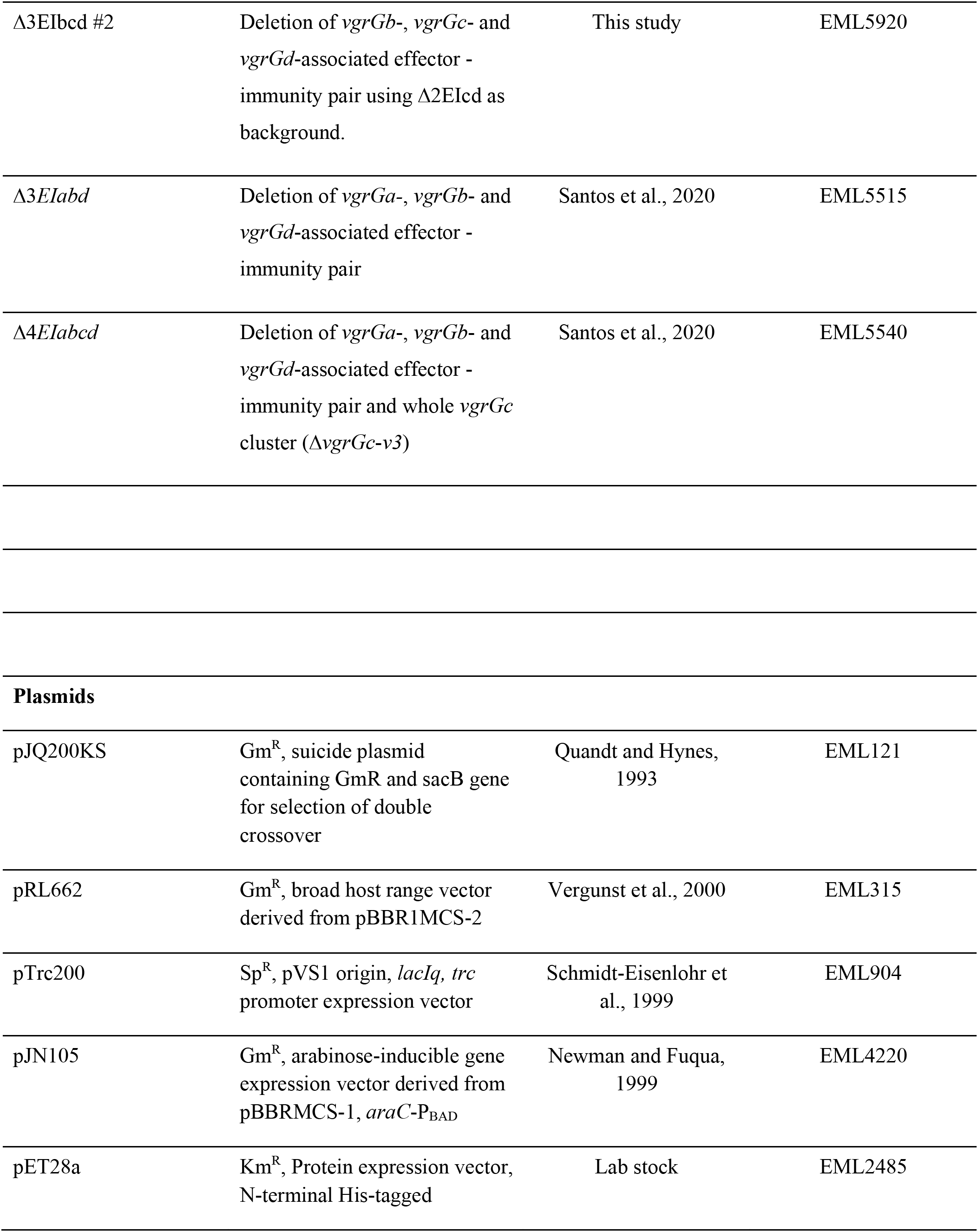
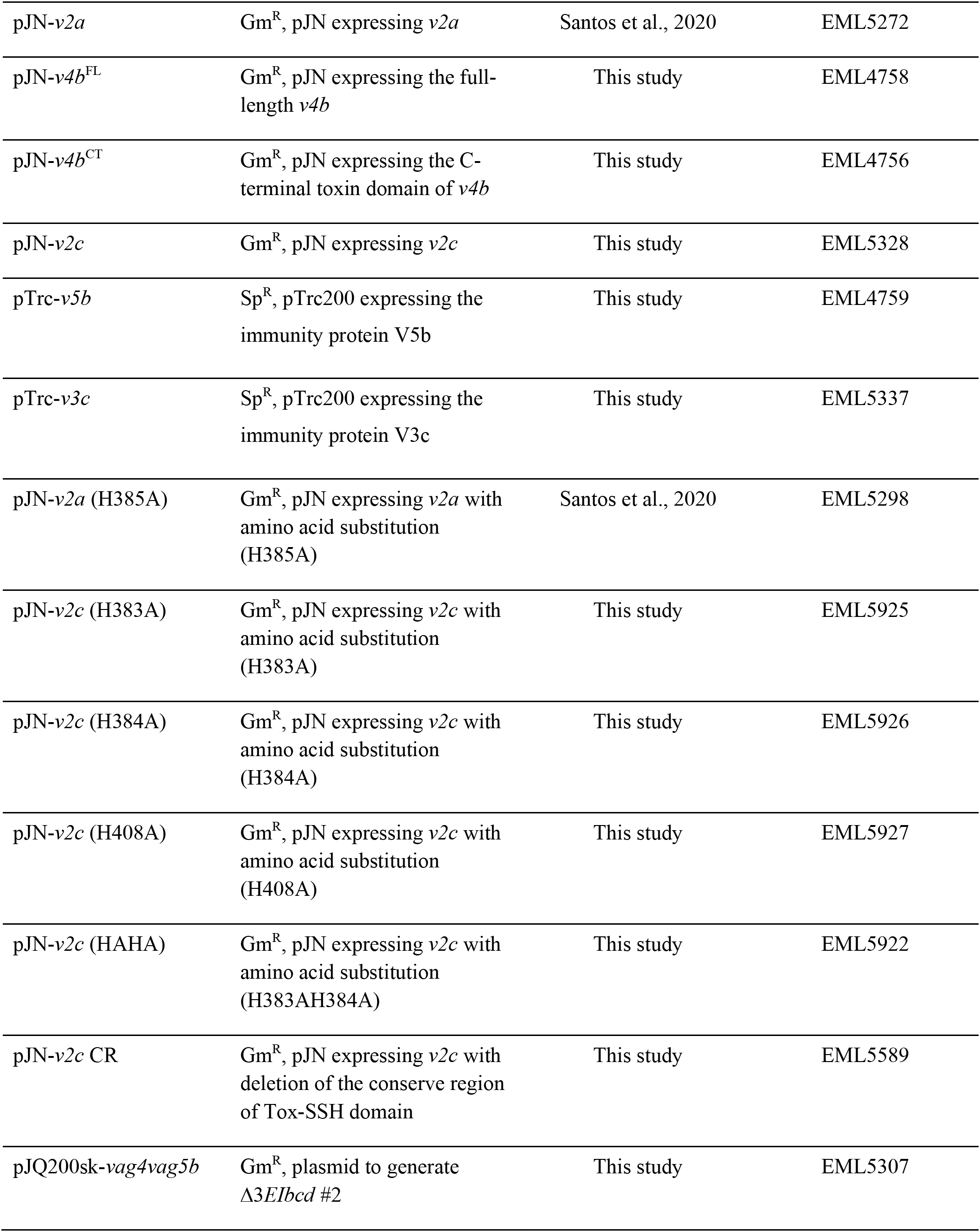
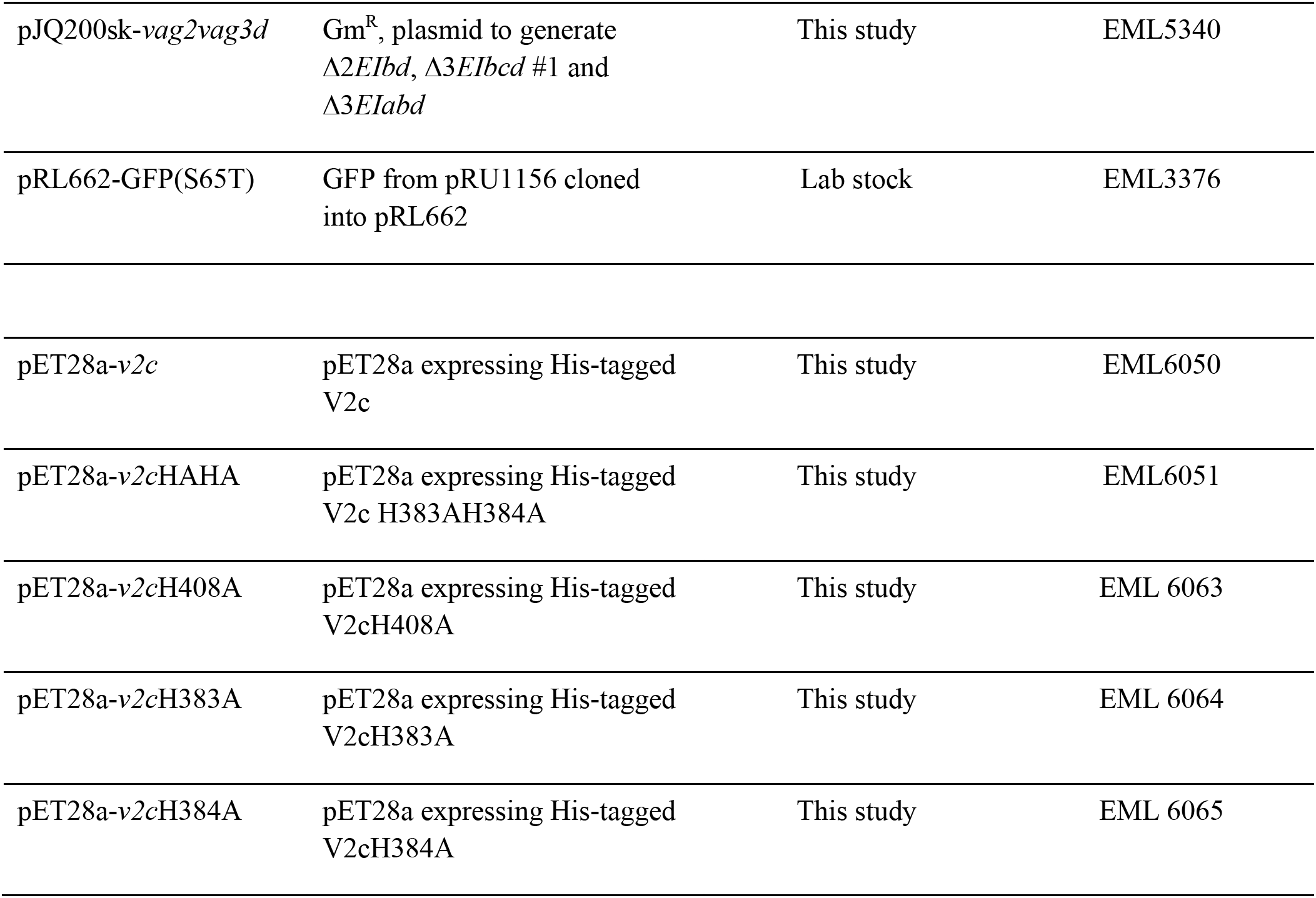
Bacterial strain and plasmids used in this study.

**Supplementary Table S2.**
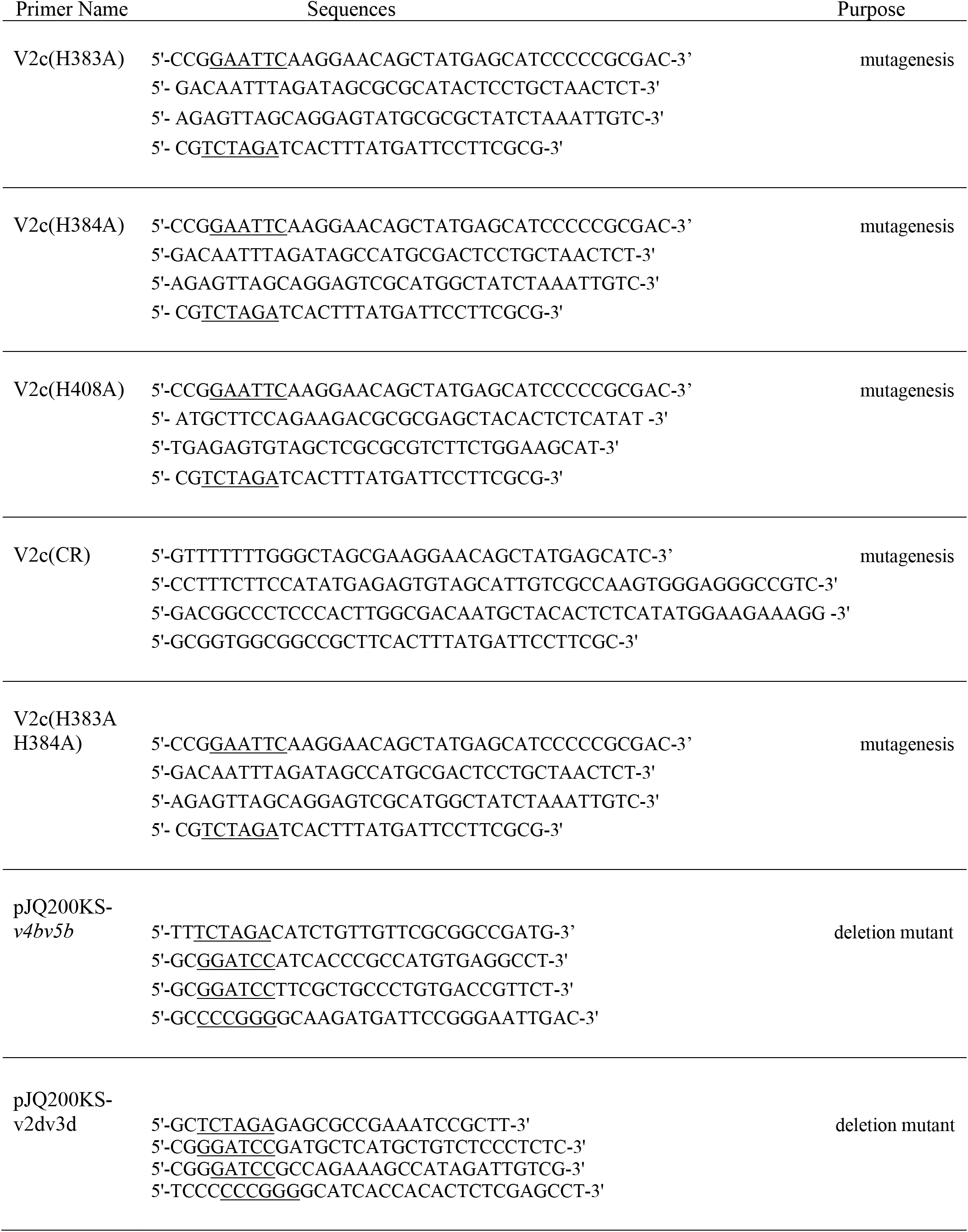
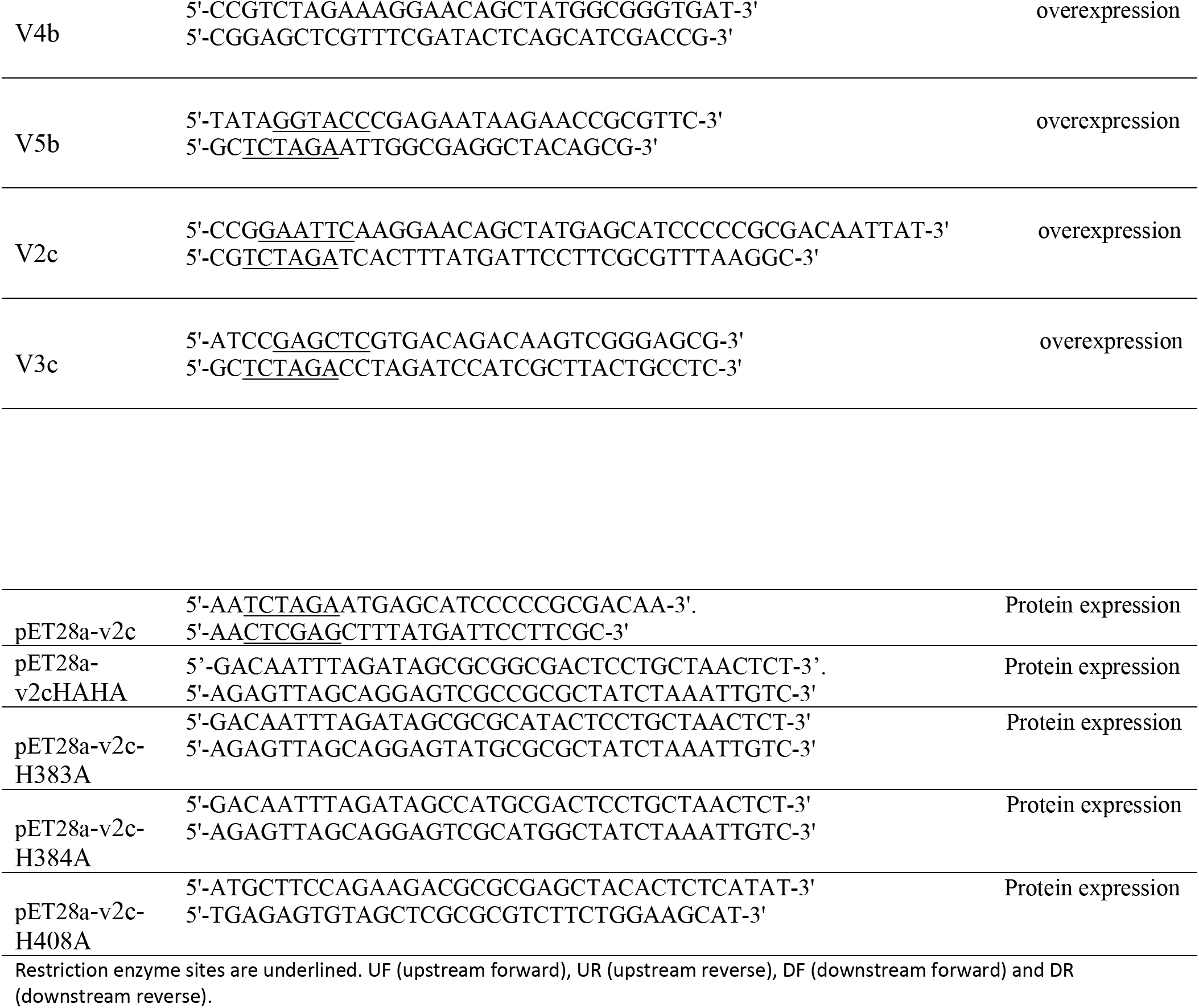
Primers used in this study.

